# Fine Tuning Genetic Circuits via Host Context and RBS Modulation

**DOI:** 10.1101/2024.07.20.604438

**Authors:** Dennis Tin Chat Chan, Lena Winter, Johan Bjerg, Stina Krsmanovic, Geoff S. Baldwin, Hans C. Bernstein

**Affiliations:** Faculty of Biosciences, Fisheries and Economics, UiT - The Arctic University of Norway, 9019, Tromsø, Norway; The Arctic Centre for Sustainable Energy, UiT - The Arctic University of Norway, 9019, Tromsø, Norway; Department of Life Sciences, Imperial College London, South Kensington, London SW7 2AZ, UK; Imperial College Centre for Synthetic Biology, Imperial College London, South Kensington, London SW7 2AZ, UK

**Keywords:** Synthetic Biology, Toggle Switch, Chassis-Effect, Context Dependence, Biodesign, RBS, Genetic Circuit, Modularity, Stutzerimonas, Non-model, Host Organism, Host-Circuit Interaction

## Abstract

The choice of organism to host a genetic circuit – the chassis – is often defaulted to model organisms due to their amenability. The chassis-design space has therefore remained underexplored as an engineering variable. In this work, we explored the design space of a genetic toggle switch through variations in nine ribosome binding sites compositions and three host contexts, creating 27 circuit variants. Characterization of performance metrics in terms of toggle switch output and host growth dynamics unveils a spectrum of performance profiles from our circuit library. We find that changes in host-context causes large shifts in overall performance, while modulating ribosome binding sites leads to more incremental changes. We find that a combined ribosome binding site and host-context modulation approach can be used to fine tune the properties of a toggle switch according to user-defined specifications, such as towards greater signaling strength, inducer sensitivity or both. Other auxiliary properties, such as inducer tolerance, are also exclusively accessed through changes in host-context. We demonstrate here that exploration of the chassis-design space can offer significant value, reconceptualizing the chassis-organism as an important part in the synthetic biologist’s toolbox with important implications for the field of synthetic biology.

**GRAPHICAL ABSTRACT:** 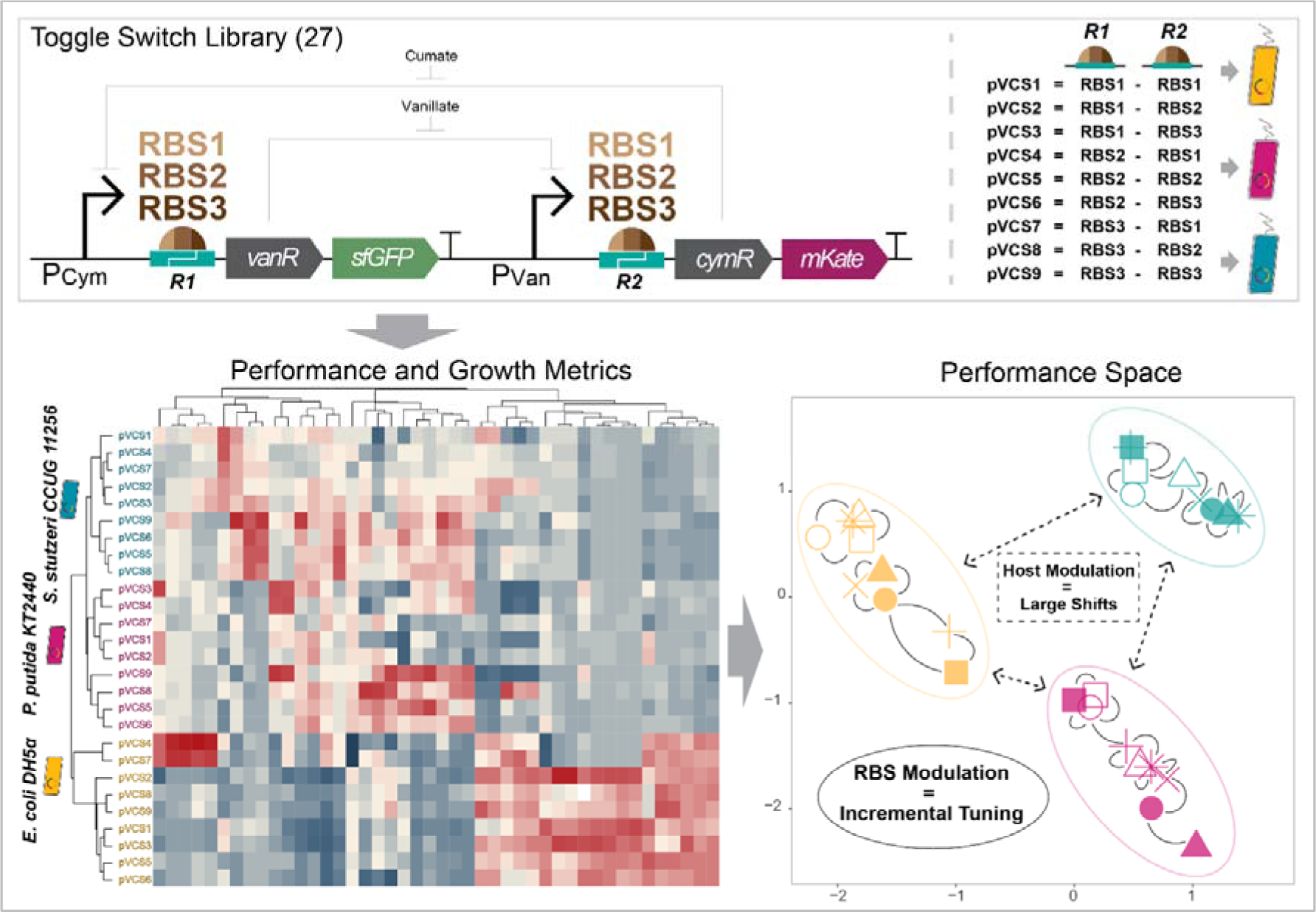

## INTRODUCTION

Genetic circuits have emerged as powerful tools for engineering cellular behavior^1^, thus opening new opportunity for biotechnology to address pressing issues such as disease mitigation^2^, climate change^3^ and food security^4^. Realizing these promises requires synthetic biologists to enhance the reliability and scope with which biological systems can be engineered to achieve desired performance characteristics. The systematic Design-Build-Test (DBT) engineering framework and implementation of standardization^5^ infrastructure within various aspect of synthetic biology has significantly accelerated advancements in this field. In the engineering of genetic circuity, the DBT cycle is typically implemented through two primary approaches: rational forward engineering and combinatorial engineering. Forward engineering involves designing high-level constructs based on available and characterized parts, relying heavily on existing knowledge of these basal components and their interactions^6^. Meanwhile, combinatorial engineering approaches attempt to exhaust large design spaces and, in some cases, all possible combinations of basal parts followed by downstream screening for desired performances^7^. Contemporary biodesign often integrate these two approaches, using forward engineering to establish the core logic and structure of genetic circuits, while combinatorial methods are employed to fine-tune performance specifications through modular component assembly^6^. This hybrid strategy maximizes the efficiency and efficacy of genetic circuit design^8^.

The performance and predictability of synthetic biological devices are influenced by the context in which they operate from^9,10^. Interactions between the host cell – i.e., “chassis” – and heterologous circuit machinery is complex and necessitate careful consideration to effectively navigate and exploit the genetic circuit performance space to attain desired attributes^11^. Addressing these challenges requires precise tuning of gene expression within synthetic circuits, which has previously been achieved by modulating regulatory elements – or “parts” – in a combinatorial manner. This includes elements such as promoters^12,13^, copy numbers^14,15^, terminators^16^ and ribosome binding sites (RBSs)^17,18^ designed in various circuit topologies^19,20^. Contemporary combinatorial engineering efforts have successfully enhanced yields of microbial cell factories^14,15,21^ and improved fold-induction of biosensing devices^22,23^ by perturbing genetic components. Tuning gene expression through RBS modulation has been a particularly popular engineering strategy for multiple reasons^24–26^. For example, the relatively short length of RBSs, which often includes spacer regions and upstream 5’-UTR components, constrains the design space to a manageable size and cost-efficient level^27^. Still, a single nucleotide change can lead to significant differences in translational strengths^26,28^ meaning a wide spectrum of gene expression levels can be achieved. RBSs and ribosome structures are also highly conserved across prokaryotes, thereby posing lower risk of unspecific interactions, compared to promoter parts which are susceptible to cross talk with transcription factors^17^. Lastly, the development of tools such as the RBS calculator by Salis (2011) to predict translation initiation rates from RBS sequence alone has further improved the efficiency of RBS modulation^29,30^.

While significant progress has been made in manipulating genetic elements, the “chassis-design space” remains relatively under explored^11,31,32^. The choice of chassis to host engineered genetic circuits usually defaults to a genetically tractable model organism (e.g., *Escherichia coli*) despite the model organism not necessarily being the most optimal host^33^. This has significant implications because the choice of host can dramatically influence circuit performance^34,35^, a phenomenon known as the chassis-effect. The chassis-effect arises from host-circuit interactions, the coupling of heterologous circuitry to endogenous cellular biology and the complex interplay between resource competition^36,37^ and potential regulatory cross talk such as promiscuous transcriptional factors^38^. The chassis-effect presents both challenges and opportunities for biodesign. While unspecific interactions and resource competition between a synthetic circuit and the host’s native genetic machinery can undermine predictability and select against circuit-bearing cells, strategic exploitation of the chassis-effect can enhance circuit performance based on specific user-defined goals^39,40^. Adding chassis-design spaces to the arsenal of a synthetic biologist’s toolbox can revolutionize microbial biodesign strategies by enabling the discovery of performance spaces beyond the constraining reliance on traditional model organisms. Taking advantage of innate pragmatic phenotypes in non-traditional hosts that complements the novel function also serves as an efficient design strategy for synthetic biology^41^. Exploration of the chassis-design space has been hindered by the fact that many organisms are not amenable to genetic transformation, for reasons that are not always clear, but has been traced to incompatible origins of replications for plasmid-based devices, toxicity of heterologous products, debilitating growth burden imposed by the circuit, unsuitable transformation methods and/or uncharacterized host immune systems^42,43^.

In this study, we explored the design space of a genetic toggle switch circuit spanning nine combinations of RBS variants and successfully introduced the circuits into three host contexts (*Escherichia coli* DH5α, *Pseudomonas putida* KT2440 and *Stutzerimonas stutzeri* CCUG11256), creating a library of unique toggle switch variants. We systematically demonstrate the chassis-effect within this library, identifying key principles for the combinatorial use of RBS and chassis-contexts to explore a tunable design space. Our findings underscore the potential of integrating RBS modulation and host-context variation to shape the performance landscape of genetic circuits, providing valuable insights for the broader field of synthetic biology.

## RESULTS

### Establishing the design space via combinatorial hosts and RBS sequences

We identified a unique performance landscape from three bacterial hosts operating a suite of genetic toggle switches constrained to the same design space defined by combinatorial pairings of RBS parts. We constructed the pVCS plasmids, a series of nine toggle switches with modulated combinations of RBS strengths regulating for the translation of the two genes coding for repressive transcriptional factors (Fig. 1a). The pVCS series was assembled using the DNA-BOT^44^ platform via automated BASIC DNA assembly^45,46^, using RBSs of known translational strengths (RBS1, RBS2 and RBS3, in increasing strength) incorporated into the BASIC linkers. The core design of our toggle switches draws inspiration from the canonical Gardner et al. (2000) genetic toggle switch, consisting of two antagonistic expression cassettes to create a bistable motif with each cassette being regulated by a negatively inducible promoter. Transcription from either promoter leads to production of the opposite inducible promoter’s cognate repressor protein and a unique fluorescent protein, thereby establishing a mutual inhibitory regulatory network. The addition of inducers cumate (cym) or vanillate (van) positively biases transcription from the P_Cym_ and P_Van_ promoters, respectively. Besides the RBS parts, the intergenic context was kept constant across all toggle switches (Fig. 1b). This includes the 5’-UTR and spacer region upstream of the start codon for the modulated RBSs and the RBSs regulating for the two genes encoding fluorescent reporters (UTR3-RBS3) (Fig. 1a).

**Figure 1.**
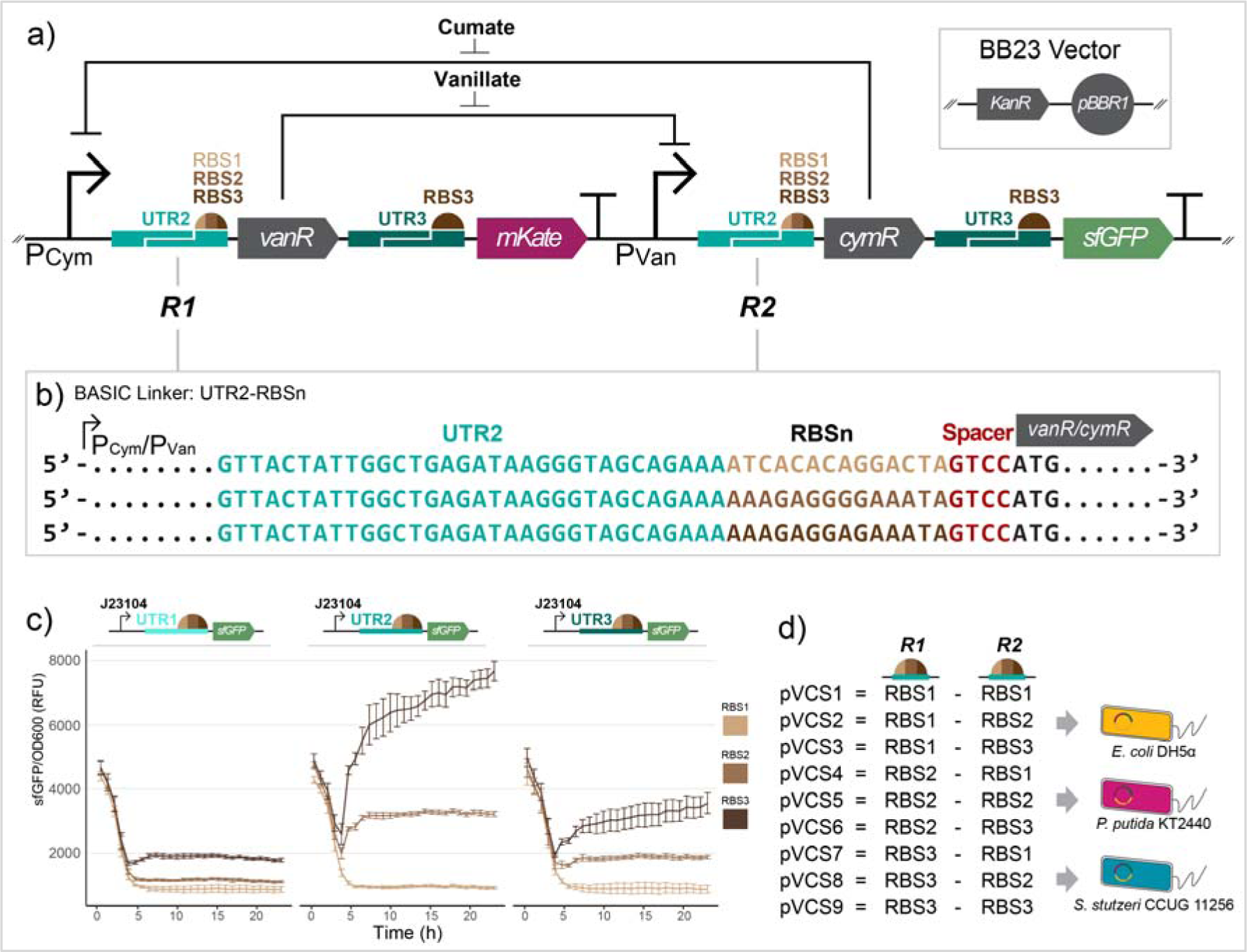
The nine pVCS toggle switches with combinatorial RBS strengths were introduced into three bacterial host contexts. a) Core design of the cumate and vanillate inducible toggle switches, varying in the combination of RBS parts upstream of genes encoding for the two transcription factors (site *R1* and *R2*). Toggle switches were cloned into the BB23 vector with *KanR* selection marker and pBBR1 origin of replication, yielding the pVCS plasmid series. b) Besides the RBS parts regulating for the transcription factors, the intergenic context was held constant across designs via the BASIC assembly linker, with each RBS being preceded by the same 5’-UTR sequence (UTR2) and downstream spacer region. RBS strength regulating for reporter protein CDSs was preceded by UTR3. c) Fluorescence assay of constitutive reporter circuits designed with all nine BASIC RBS spanning three 5’-UTR regions and three RBS parts. J23104 is a constitutive Anderson promoter. Error bars show standard deviation, n = 4. d) The pVCS series of nine toggle switches were successfully introduced into three host contexts, *Escherichia coli* DH5α, *Pseudomonas putida* KT2440 and *Stutzerimonas stutzeri* CCUG 11256.

A preliminary assay with constitutive fluorescence reporter constructs verified the variable translational strengths of the three RBS parts (Fig. 1c) used in this study. The assay also reveals how upstream 5’-UTR identity can greatly impact gene expression levels, the effect of which is most prominently observed for the RBS3 linker set, with a six-fold difference in estimated steady-state fluorescence levels between UTR1-RBS3 (1860 ± 50 RFU) and UTR2-RBS3 (7010 ± 270 RFU) circuit variants. Open-Source Translation Initiation Rate (OSTIR) program also infers the expected increasing translation initiation rate according to the predetermined strengths of the RBS parts under the context of the actual toggle switch designs (Supplementary Figure S1). Together, these results lend power to the use of the BASIC RBS linkers as a strategy to fine-tune circuit performance The pVCS plasmid series, using the pBBR1 origin of replication, was transformed successfully into all three host species (Fig. 1d), yielding a library of 27 toggle switches spanning nine RBS combinations and three host contexts. With this experimental framework we set out to characterize the performance variability under standardized conditions and elucidate the efficacy of the two design spaces as strategies to tune circuit function.

### Variable RBS pairings lead to diverse performances across hosts

Comparison of performance profiles among toggle switch variants reveals that host-context has a more significant influence than RBS-context on device performance. Specifically, variations in host-context lead to more substantial shifts in the overall performance profile, whereas changes in RBS-context result in more incremental adjustments. Performance metrics were derived from the fluorescent response dynamics of the circuit variants across induction states in a toggling assay (Fig. 2a). These measurements include measurements of lag time (Lag, in units of h), rate of exponential fluorescence increase (Rate, RFU/h), and the steady-state fluorescence output at stationary phase (Fss, in units of RFU). The biological interpretations of these metrics vary according to the state of induction. For instance, in absence of inducer, the Rate and Fss metrics for sfGFP and mKate represent expression leakage or baseline output. Conversely, sfGFP output in presence of cym (or mKate in presence of van) indicate expression leakage from the P_Van_ promoter despite supposed VanR repression. We also define fold-induction (FI) as the ratio between induced and non-induced Fss, which informs of the responsive range of the toggle switch from baseline reference, to induced signal.

**Figure 2.**
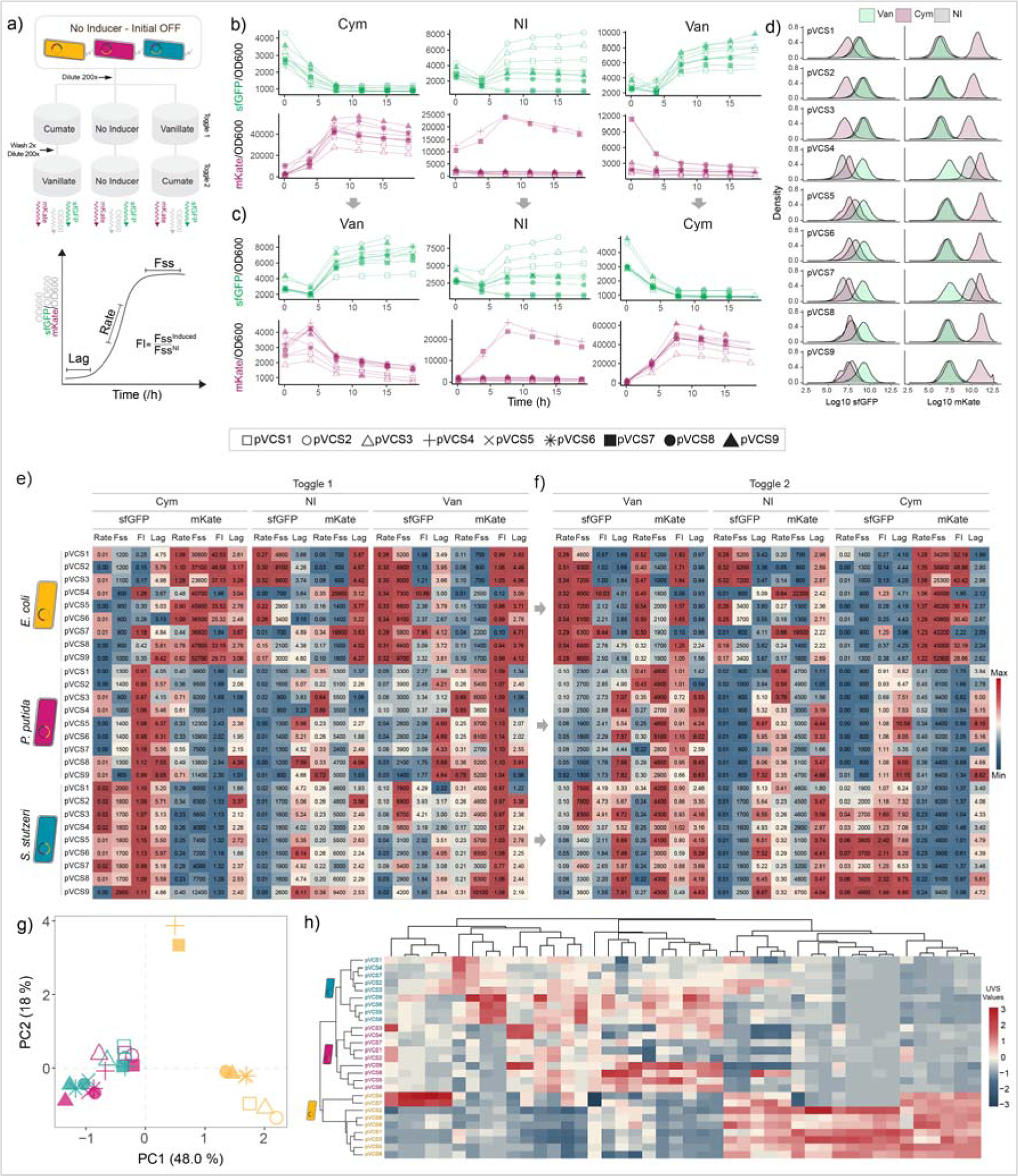
Toggle switch performance clusters by similar host context rather than RBS context. a) Schematic of toggling assay. Cells in initial OFF state were diluted to 0.75 mM cym and van, as well as a no inducer (NI) control. To toggle, cells were washed twice before being diluted 200x to respective opposite induction state under same concentration. The metrics Lag (lag time before fluorescence increase in units of hours), Rate (rate of exponential fluorescence increase in units of RFU/h) and Fss (steady-state fluorescence at late phase in units of RFU) were estimated from growth normalized sfGFP and mKate fluorescence curves. Fold-induction (FI) is defined as the ratio between induced and non-induced Fss. b) Representative growth normalized fluorescence dynamics across induction states for toggle switch-carrying *E. coli* cells toggled from initial OFF and c) toggled to opposite induction state. d) sfGFP and mKate fluorescence intensity distribution of *E. coli* cell populations. Performance metrics across the 27 toggle switch variants from e) Toggle 1 and f) Toggle 2 across induction state for both outputs. Color scale indicates maximum and minimum value relative to each column. g) Principal component analysis plot of all quantified toggle assay metrics. h) Euclidean distance-based hierarchal clustering of circuit contexts and toggle switch performance metrics. UVS = unit-variance scaled. n = 4.

In toggle 1, cells were induced from an initial-OFF state to van-ON or cym-ON while also maintaining a control culture with no inducer (NI). A second phase of the circuit, toggle 2, was then activated with the alternate induction state after dilution and subsequent growth to stationary phase (Fig. 2b, c, Supplementary Figure S2). Examining the fluorescence responses at individual cell level via flow cytometry verified a uniform population response from all 27 toggle switch variants (Fig. 2d, Supplementary Figure S2). Comparing the quantified performance metrics reveals a diverse range of performance profiles, demonstrating how varying RBS and host context can grant access to a wider set of performance specifications (Fig. 2e). Switches operating within the context of *E. coli* exhibited an overall stronger sfGFP and mKate response, as indicated by the higher Rate and Fss values. For instance, cym-induced mKate Fss levels range from 23600 ± 1500 RFU (pVCS3) to 52700 ± 1200 RFU (pVCS9) in *E. coli*, while the highest mKate Fss values achieved in *P. putida* and *S. stutzeri* were 13800 ± 500 (pVCS8) and 12400 ± 170 RFU (pVCS9), respectively, two-fold lower than the lowest output attained by pVCS3 in *E. coli*. This suggests that *E. coli* provides an environment conducive to higher fluorescent protein accumulation.

A high fold-induction (induced signal to baseline ratio) is usually desired in genetic devices such as chemical event detectors^34,47^. When implementing this current toggle switch design into *E. coli*, the circuits gain access to much higher fold-induction response upon cym induction (measured via mKate fluorescence reporting) as compared to van measured by sfGFP. While *E. coli* cells exhibit some of the highest sfGFP Fss values among all circuits, they were all accommodated with high baseline, resulting in overall similar responsiveness across hosts (average sfGFP_Van_ FI value for all twenty-seven strains is 3.0 ± 2.1). Accessing circuit variants that operate similarly in function but at different amplitudes can be beneficial, especially when the baseline signal of some circuits saturates the measurable range of a fluorescent reader or when background noise in samples is high.

Cells toggled between induction states showed the expected inversion of fluorescence response according to their design (Fig. 2f), albeit with some devices toggled from van-ON to cym-ON exhibiting a slight attenuated response in *P. putida* and *S. stutzeri*. This response attenuation appears to be dependent on past induction states, as cells toggled from cym-On to van-ON demonstrate similar performance as cells toggled from initial-OFF across contexts and only minute concentrations should remain after washing and dilution. Principal Component Analysis (PCA) of performance profiles dataset clusters toggle switches in *P. putida* and *S. stutzeri* into their own cluster with almost equal dissimilarity to *E. coli* (Fig. 2g). The spread of toggle switches within each host-cluster illustrates the incremental adjustments that occur when varying RBS combinations, highlighting the fine-tuning capability of varying RBS parts. Notably, *E. coli* pVCS4 and pVCS7 forms their own distinct cluster, diverging in behavior even from other *E. coli* toggle switches. Indeed, closer inspection of pVCS4 and pVCS7 metrics reveals a high mKate and low sfGFP baseline, suggesting the native state of these designs are more biased towards expression from the P_Van_ promoter, likely due to their unique RBS combination. This opposite base-state appears to be the cause of their opposite behavior. For instance, the mKate_Cym_ FI values of these two switches are 2.0 ± 0.1 and 1.8 ± 0.1 respectively, manifolds lower than the FI values of the other seven *E. coli* switches, which range from 25.3 ± 1.0 to 46.6 ± 2.7. Meanwhile, pVCS4 and pVCS7 achieves the highest recorded sfGFP_Van_ fold-induction values out of any device (10.9 ± 0.7 and 8.0 ± 0.5, respectively).

The different hosts revealed preferential base level operation of the toggle switch under non-induced state. Toggle switches in *P. putida* and *S. stutzeri* demonstrate higher average mKate Fss values (mKate_NI_ Fss 4600 ± 1000 RFU and 5200 ± 1900 RFU, respectively) than those in *E. coli* (average 1100 ± 500 RFU among circuits, excluding pVCS4 and pVCS7). The opposite is observed for sfGFP baseline output, with toggle switches in *E. coli* exhibiting higher average sfGFP_NI_ Fss values. These observations suggest that the host environments greatly affect the native toggled state of the device, which in turn affects circuit performance and underscores the substantial impact of host context on genetic circuit behavior.

Hierarchical clustering of performance profiles reveals that toggle switches separate according to the three host contexts, branching *P. putida* and *S. stutzeri* into their own clusters (Fig. 2h). Interestingly, within the *P. putida* and *S. stutzeri* cluster, specific combinations of RBS strengths found in pVCS1, pVCS2, pVCS3, pVCS4 and pVCS7 cluster separately from the other four circuits. A similar clustering within *P. putida* and *S. stutzeri* clusters is observed in Fig. 2g, indicating in cases of little difference between hosts, similarity in RBS combinations does leads to similar performance. Overall, our results suggest that changes in host context cause significant shifts in overall toggle switch performance, whereas modifications in RBS context typically results in minor changes. However, in certain instances, such as with the pVCS4 and pVCS7 variants in *E. coli*, RBS changes can also greatly alter circuit performance, highlighting the important of exploring a wide design space.

### Species-specific growth physiology imposes limits on toggle switch performances

Introduction of a genetic circuit into a host environment couples the exogenous circuit to the host’s cellular machinery, causing a reallocation resourceless originally dedicated to growth and cell maintenance^36,48,49^. Circuit performance is therefore constrained by the unique growth physiology of each host and physiological responses to the heterologous toggle switch machinery^35^. To elucidate this host-device interplay on growth, we determined the growth burden caused by toggle switch operation. Growth burden was quantified through the Δμ% metric, defined as the relative percentage difference in specific growth rate (μ) between two conditions. Growth dynamics of the toggle assay were measured simultaneously and by comparing plasmid-bearing strains in baseline state against their wildtype (WT) genotype counterpart (Fig. 3a, NI-pVCS vs. NI-WT). The results show that maintaining the toggle switch consistently imposes a reduction of growth across all hosts, but to various degrees.

**Figure 3.**
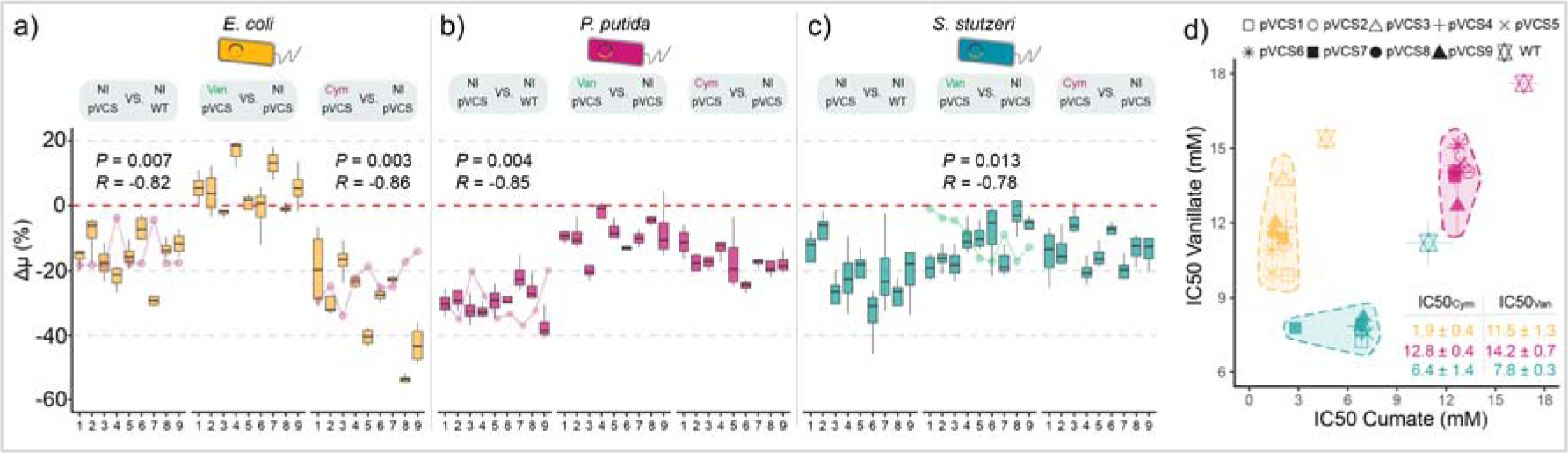
Coupling between circuit function and chassis growth leads to operational limits of toggle switches. Growth burden quantified through the relative percentage difference in growth rate (Δμ%) between non-induced (NI) plasmid-bearing strains and WT, van induced against NI and cym induced and (NI) strains for a) *E. coli*, b) *P. putida* and c) *S. stutzeri*. In comparisons between growth burden and sfGFP/mKate output metric (product of Rate and Fss) was significantly correlated, the associated *P*-value (*P*) and Pearsons’s correlation coefficient (*R*) is shown. d) Half-minimum inhibitory concentration (IC50) of vanillate and cumate for each strain. Shaded areas are clustered by host, excluding WT. Table inset shows average IC50 values within each inducer-host group. Error bars shows standard deviation, n = 4.

The most severe and consistent growth burden was observed among *P. putida* strainş with plasmid bearing cells experiencing on average −29.3 ± 4.4 % lower growth rate than their WT counterpart. *E. coli* and *S. stutzeri* cells experienced on average more moderate −15.6 ± 6.9 % and −22.2 ± 7.4 % lower growth rate compared to their WT counterparts. The pBBR1 origin of replication has been reported as a low-copy number plasmid in *E. coli*, but previous studies has shown that plasmid copy number is subject to the chassis-effect and that *P. putida* can maintain a 10-fold higher plasmid copy number of a pBBR1 plasmid compared to *E. coli*^35,50^, which could explain the higher degree of growth burden. Further induction of the switches exaggerates growth burden, with toggle switches operating from *E. coli* experiencing the most drastic growth inhibition upon cumate induction, but the degree of growth burden ranges widely from −16.8 ± 5.4 % (pVCS3) to −53.0 ± 1.8 % (pVCS8). Pearson correlation analysis reveals significant negative correlation between mKate_Cym_ output and Δμ% metric in *E. coli* variants induced with cym (*P*-value = 0.007, *R* = −0.82). This suggests the degree of growth burden correlates with output, which is in turn tunable with RBS. The same negative significant trend is also observed for van induced *S. stutzeri* variants and sfGFP_Van_ output and non-induced mKate output in *E. coli* and *P. putida*. Stronger output thereby leads to greater growth burden, which has major implications towards tuning genetic circuits within the bounds of species-specific growth constraints. This result also highlights the importance of optimizing not only for performance, but also for growth when it comes to working with living systems. We have previously demonstrated that differences in growth physiology significantly correlated with differences in performance^35^, which we once observe when comparing the differential performance of all 27 toggle switches and their differential growth physiologies through Procrustes Superimposition analysis, solidifying our previous findings (Supplementary Figure S3). Exploring both host and RBS design space allows the discovery of a suitable version of a circuit that balances growth and performance.

The results shown thus far establish an interplay between device and growth physiology, which is complex and depends on the specific choices of bacterial host species, RBS combinations and user-defined operation of the toggle switch. Our results show clearly that toggle switches result in burdened growth and therefore tax the host of its cellular resources. This effect results in lower cell division rates, which in turn impose reciprocal constraint back onto circuit performance within the operational concentration limits defined by the induction kinetics of P_van_ and P_cym_, respectively. We identified these limits by determining the half-minimal inhibitory concentration (IC50) of each inducer module (Fig. 3d). A clustering of toggle switches by their host context is again observed. *P. putida*, an established model organism known for its robustness^51^, is the most tolerant to both inducers with some of the highest IC50 values, particularly for cumate. *E. coli* toggle switches exhibit the lowest tolerance to cumate, with an average IC50_Cym_ of 1.9 ± 0.4 mM. WT strains of hosts all demonstrate higher tolerance to inducers, as expected due to not being burdened with maintaining a foreign plasmid. The clustering of IC50 values within each inducer-host group shows minimal variation among RBS variants, highlighting the limitations of using RBS modulation to enhance inducer tolerance. In contrast, altering the host context emerges as an effective method to enhance inducer tolerance and by extension can be used to engineer forth performance capabilities beyond those achievable through intra-circuit context alone.

### The RBS design space allows for precision tuning of species-specific operation objectives

Modulating host-context was found to greatly alter the induction kinetics of the genetic circuits, whereas varying RBS strength is ideal for fine-tuning of performance objectives. The induction kinetics of all 27 toggle switch variants were determined to gain a comprehensive assessment of their performance across a wider range of concentration states (Fig. 4a - f), yielding deeper insights into the unique performance characteristics not revealed from a single induction point. The Hill function was fitted to induction curves to estimate the system parameters β (Fig. 4g), which represents the max fluorescence output at saturating inducer concentrations; the activation coefficient *K*, indicating the inducer concentration at which half-maximum output is achieved; and *n*, the Hill coefficient serving as a fitting parameter. The system fold-induction (FI^S^) is calculated as the ratio between β and *C*, where *C* represents the empirically determined fluorescence output at 0 inducer concentration. Collectively, these performance metrics provide a detailed description of each toggle switch variant’s performance specification.

**Figure 4.**
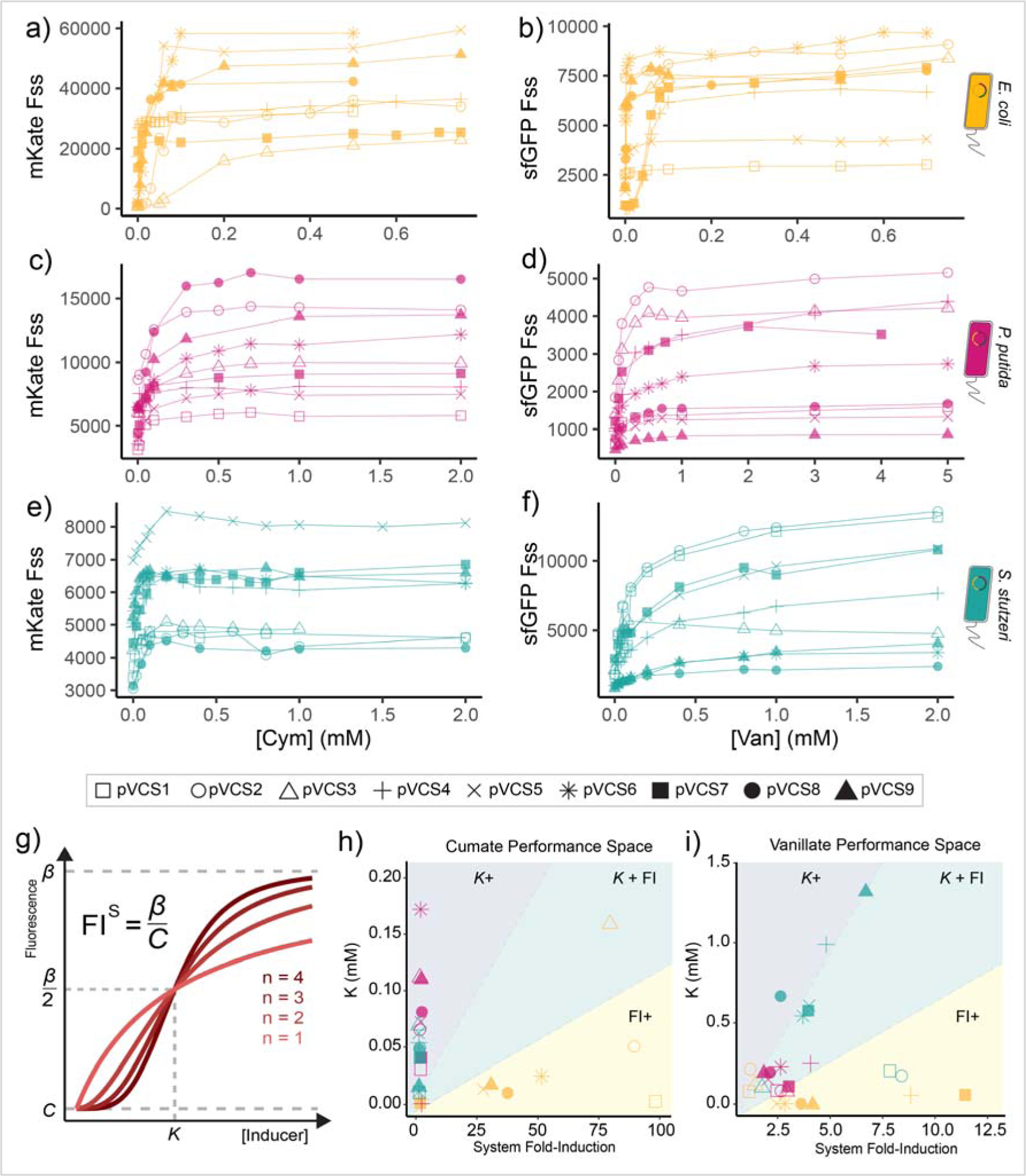
Exploration of the chassis-RBS design space reveals chassis exclusive performance spaces. Cumate and vanillate induction response curves of the pVCS toggle switches for a-b) *E. coli*, c-d) *P. putida* and e-f) *S. stutzeri*. Note differences in axis scales. g) Visual representation of Hill function curve with Hill coefficients (*n*) and the other three estimated parameters: *C* (fluorescence steady-state output in absence of inducer or leakage), *K* (activation coefficient or inducer concentration in which half-maximal output is attained) and β (fluorescence steady-state at saturating inducer levels). System fold-induction (FI^S^) is the ratio of estimated β and *C*. The sampled h) cumate and i) vanillate toggle switch performance parameter space, plotting activation coefficient and system fold-induction. Dashed lines arbitrarily divide regions designated as sensitivity optimized (*K*+), reporting optimized (FI^S^+) and both (*K* + FI^S^). Error bars how standard deviation, n = 7.

The estimated *K* values indicate that appreciable cumate and vanillate induction can occur in the micromolar concentration range, consistent with previous literature describing P_Cym_ and P_Van_ induction systems^52–55^. A wide range of inducer sensitivities is observed. For instance, the *E. coli* toggle switches demonstrated some of the lowest *K*_Van_ values, with an average value of 0.059 ± 0.005 mM, approximately ten-fold lower than the average *K*_Van_ values for *S. stutzeri* (0.54 ± 0.16 mM), suggesting the cellular context of *E. coli* alters the toggle switch’s responsiveness to induction. Meanwhile, cumate induction appears to require lower concentrations overall, as all *K*_Cym_ values range from 0.002 ± 0.004 mM to 0.156 ± 0.004 mM across all variants.

The observed performance space of the toggle switch library, visualized by plotting *K* against FI^S^ (Fig. 4h, i) reveals that certain performance specifications can only be accessed by varying host context. Considering cumate induction kinetics, toggle switches operating within *P. putida* or *S. stutzeri* context achieve FI^S^ values up to 3 at most (Fig. 4h). When operating within *E. coli* however, a toggle switch with FI^S^_Cym_ value up to 98.6 can be achieved, a clear example of how varying host-context rather than intra-genic context can be used to tune function. These *E. coli* switches however, show little variation in terms of *K*_Cym_ values, limiting their sensitivity and inducible range. Meanwhile the same toggle switches functioning within *P. putida* and *S. stutzeri* hosts (with low FI^S^_Cym_) outperform *E. coli* variants in terms of *K*_Cym_ values, exhibiting a much wider range of *K*_Cym_ values. Next, considering vanillate induction kinetics (Fig. 4i), *E. coli* switches again demonstrate relatively constant K_Van_ values across RBS variants and vary more in terms of fold-induction. *P. putida* switches form a tight cluster, showing little diversity in their performance compared to the more spread *S. stutzeri* group. Notably, varying RBS composition leads to host-specific changes in performance. For instance, switches in *E. coli* only spread along the x-axis, while *P. putida* and *S. stutzeri* variants cluster along the y-axis. The spread of toggle switch variants along each host-group suggests RBS can be used to fine-tune performance, but on the other hand, this is an example of how host-context can confine the performance of the genetic circuit. Overall, our results show that a combined approach of intergenic and inter-chassis exploration serves as a method for surveying performance spaces to gain access to circuits with more diverse performance specifications.

## DISCUSSION

Synthetic biology is steadily advancing past its proof-of-concept stage. To fully realize its full potential, the field must not only be concerned with developing proof-of-concept of novel capabilities, but also optimizing them for practical use. Advancing beyond the constraints imposed by working with the few preferred model organisms is an important step in this regard. In this work, we defined a combinatorial toggle switch design space based on RBS and host-context and observed variable device performances within this framework. The three host-contexts sampled were each associated with optimized performance. While this optimization can come with trade-offs, it showcases the potential of modulating chassis to tune circuit performance. We demonstrate that through combined adjustment of RBS strength and host-context, the performance of a genetic toggle switch can be optimized towards high sensitivity (i.e., low *K*), high induction range (i.e., high *K*) or reporting efficiency (i.e., high FI). We thereby provide synthetic biologists new insight into how the chassis and RBS combinatorial design space can be exploited to optimize genetic circuits to achieve desired outcomes. Furthermore, we show that certain parameters, such as increased inducer tolerance, are exclusively accessed by varying chassis-contexts, further highlighting the value of broadening the available chassis-design space.

Our characterization of toggle switch performance shows that the cellular environment imposes a general floor (fully repressed output) and ceiling (fully induced output) limits upon the toggle switch as well as influencing the unbiased toggled state of the circuit (expression more toggled towards P_Van_ or P_Cym_ in absence of no inducer). The influence of host-context on circuit performance can originate from a range of host-circuit interactions. For instance, differences in minimum output could be due to the promoters recruiting RNA polymerases to different degrees of efficiency (host-specific promoter strengths)^56^, differing plasmid copy number^35^, or promiscuous binding of the circuit’s transcriptional factors within the host genome ^57^ leading to higher steady-state leakage. A cellular environment that imposes a lower turnover rate on the fluorescent proteins, more efficient folding of heterologous proteins, or higher free ribosome and/or RNA polymerase could lead to increased gene expression as well. Previous studies on broad-host-range operation of genetic toggle switches have established significant correlation between differential performance and differential growth dynamics^35,58^. This can be rationalized by the fact that changes in growth represents a change in physiological state of the cell including resource pool and cell-wide parameters such as transcription, translation, degradation and dilution rates^59^. Procrustes Superimposition analysis on our expanded sample size of 27 toggle switches reports that variants with more (dis)similar growth dynamics also exhibit more (dis)similar performance. Growth dynamics as determined here in this study can be practically measured compared to other gene expression parameters (e.g., ribosome or RNA polymerase abundance), and its significant correlation with performance thereby makes it a practical input parameter for machine learning algorithms to predict chassis-effect^60^. Integrating machine learning to predict genetic circuit performance would require greater standardization in the empirical characterization of genetic circuits^61,62^.

The practical application of a designed microbial system requires multiple performance parameters to be optimized^33^, and a top performer is often selected for by compromising between parameters^7^. For instance, as we report in this work, high output levels are known to be negatively correlated to growth^63^ (Fig. 3), the latter being a crucial factor that must be balanced for when working with living systems. Selecting for an optimal performer must be guided by user-defined specifications and mission requirements. Considering our toggle switches as example circuits, the markedly high fold-induction of most toggle switches implemented in *E. coli* makes them the most optimal cumate sensing devices, but their low cumate tolerance limits them to cases where cumate concentration does not exceed well beyond 1.9 mM. For higher concentration detection, users will have to select a toggle switch implemented within *P. putida*, which displays higher cumate tolerance, while sacrifice reporting strength within acceptable levels. Besides fold-induction and inducer tolerance, factors such as response time, signal amplitude and sensitivity must also be weighed and balanced to determine the optimal performer. The signal amplitude of the system must be within the detectable range of the measurement device available (suitable *C* and β). The chosen system must be sensitive enough to be able to detect the chemical in question (low enough *K*) and if titration of sample concentration is desired, a circuit with high *K* would be optimal (circuit in *P. putida* or *S. stutzeri*). On the other hand, if only the presence/absence of a given chemical is of concern, parameter *K* can be sacrificed for shorter response time (a smaller lag phase) and higher fold-induction (circuit in *E. coli*). Sampling a wider performance space increases the chances of discovering optimal performance specifications, and we have here exemplified the value of exploring the chassis-design space by demonstrating how host-context can be leveraged to gain access to performance specifications not achievable through RBS alone.

Numerous methods to minimize context dependency in hopes of increasing stability and predictability of genetic circuits has been developed. This includes tools to segregate interactions between heterologous and native components (i.e., orthogonalization) and the practice of genome reduction^64^. Orthogonalization includes use of orthogonal ribosomes^65,66^, orthogonal RNA polymerases^67^ and the practice of encoding circuit DNA in an alternative codes only decipherable by said orthogonal machinery^68^. A reduced genome context with only genes essential for growth in controlled laboratory conditions is believed to mitigate the risk of unspecific interactions and increased host fitness^66^. Indeed, there is a prevalent notion that an ideal chassis organism is one with a reduced genome due its supposed reduced context complexity^69–71^. In contradiction to this notion, we, and others^34,39^, have showed that there is value to be gained by instead taking advantage of the contextual diversity that resides within different chassis-organisms. Our findings push the reconceptualization of the role of the chassis-organism as both a valuable part in the synthetic biology toolkit and a design variable. This notion promotes the strategic utilization of the inherent diversity found in different species to optimize genetic circuit performance. While added complexity can certainly mean increased risk of failure, this can be overcome by expanding the explored design space through high through-put DNA assembly and screening technology, which is continuously advancing^72^. Better yet, a merging of strategies from the two perspectives, such as applying tailored genome reduction to chassis-organisms that maintains the desired innate phenotypes can reap benefits of both approaches. Examples of targeted genome deletion studies on both model and non-model organisms has already resulted in improved user-defined performance^64,73–75^. With the steady increasing number of organisms with pragmatic phenotypes being domesticated as biotechnology platforms^76–80^, we envision a future where choice of host organism becomes a staple design factor in combinatorial engineering endeavors, in equal footing of promoters and RBS strengths, which will surely enhance our ability to engineer forth solutions through biology.

## Supporting information

Supplementary Figures, Tables and Other

## ACKNOWLEDGEMENTS

We thank the SEVA repository for their donation of pSEVA231 plasmid.

## AUTHOR CONTRIBUTIONS

Conceptualization, DTCC and HCB; Methodology, DTCC and HCB; Software, DTCC; Investigation, DTCC, LW, JB and SK; Resources, DTCC and GSB; Data Curation, DTCC; Writing – Original Draft, DTCC and HCB; Writing – Review & Editing, DTCC, GSB, HCB, LW, JB and SK; Visualization, DTCC; Supervision, HCB; Funding Acquisition, HCB.

## DATA AND CODE AVAILABILITY

Experimental data files and R MarkDown scripts used for analysis and plotting are publicly available online on the Open Science Framework database as part of the project name *Chan.RBS.Host.Context* (https://osf.io/4ye9w/).

## MATERIALS AND METHODS

### Species, cultivation, cloning, and transformation

Overview of species used in this study can be found in Supplementary Table S1. Cells were cultured in LB media at 37 °C unless specified otherwise, inoculating with single streaked colonies. BB23 backbone and pVCS-carrying strains were cultivated in presence of 50 µg/mL kanamycin while wild types were grown without. 199 µL of media was inoculated with 1 µL of overnight culture in black clear-bottom 96-well plates (Thermo Fischer, 165305) and sealed with Breath-Easy film (Sigma-Aldrich, Z380059). OD_600_, sfGFP (Ex 485/ Em 515, gain 75) and mKate (Ex 585/ Em 615, gain 125) fluorescence was measured continuously using a Synergy H1 plate reader (Agilent Biotek, Serial Number 21031715) with continuous linear shaking (1096 cpm, 1 mm) at 9 mm read height. Working stock solutions of 1 M vanillic acid (Thermo Fisher Scientific, 10228789) stock and 1 M cumate (Sigma-Aldrich, 536663) was prepared by dissolving powder in 70% ethanol respectively supplemented with 150 µL of 3 M of NaOH. Cloning was performed using *E. coli* DH5α, made chemically competent and transformed following the Inoue method^81^. *P putida* and *S. stutzeri* were transformed via electroporation method as previously described^35^.

### Automated BASIC Assembly and DNA-BOT

pVCS plasmids were assembled in the Biopart Assembly Standard for Idempotent Cloning (BASIC)^44,45^ cloning environment. Automated BASIC assembly was performed as described in Storch et al. (2019)^44,46^ using the DNA-BOT platform with the OpenTrons2 liquid handling robot (Opentrons, 999-00111) and temperature module. BsaI-HFv2 restriction enzyme and T4 DNA ligase was purchased from New England Biolabs (R3733 and M0202L, respectively). Mag-Bind TotalPure NGS (Omega Bio-Tek, M1378-01) was used to manually purify restriction–ligation reactions as per the manufacturer’s instructions. DNA sequences for the pVCS part components can be found in Supplementary Table S2. Part and assembly input files for the DNA-BOT can be found in Supplementary Material S1.

### RBS Translation Initiation Rates Prediction with OSTIR

OSTIR (Open-Source Translation Initiation Rates, v1.1.2, https://github.com/barricklab/ostir) was run with default settings, using the highly conserved anti-Shine-Dalgarno sequence of *E. coli* (5’-ACCTCCTTA-3’)^82^. Briefly, OSTIR employs a thermodynamic model (ViennaRNA^83^) of bacterial translation initiation to calculate the Gibbs free energy of ribosome binding, which infers the Gibbs free energy change to a protein coding sequence’s translation initiation rate and thereby expression strength. As input sequence, the 33 bp 5’-UTR linker region upstream of the 14 bp RBS was included. The RBS is immediately followed by a 4 bp (5’-GTCC-3’) spacer region and subsequently the gene CDS, which all begin with the start codon 5’-ATG-3’. Genes *cymR* and *vanR* have the 5’-UTR2 (5’-TGTTACTATTGGCTGAGATAAGGGTAGCAGAA-3’) sequence upstream of their RBS, while genes *sfGFP* and *mKate* have the 5’-UTR3 (5’-GTATCTCGTGGTCTGACGGTAAAATCTATTGT-3’).

### Toggle and growth assay and flow cytometry

Overnight culture grown in absence of inducer was used to inoculate media in 96-well plates supplemented with cym, van and no inducer condition. To toggle, cells were harvested by centrifugation at 4000 RPM for 20 minutes at room temperature and supernatant removed before resuspending in 200 µL LB media, this washing step was repeated for a total of two washes. After final resuspension, 1 µL of washed cells were inoculated to 199 µL fresh media supplemented with the opposite respective inducer. Growth difference metric Δµ was calculated using equation 2.

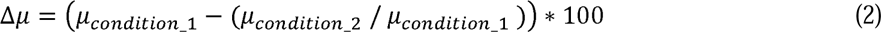

Where *µ* is max specific growth rate and “condition_1” and “condition_2” denotes sample condition in terms of genotype and induction state. In R, max rates of OD_600_ and normalized fluorescence curves were estimated based on a rolling regression method using the “all_easylinear” function from the growthrates (v.0.8.4, https://CRAN.R-project.org/package=growthrates) R package. Lag times and curve plateaus of OD_600_ and normalized fluorescence curves were determined using the “all_growthmodels” function, fitting the Gompertz growth model^84^ with additional lag (λ) parameter.

Van-ON, Cym-ON, and NI cells at exponential late phase were fixed with formaldehyde to a final concentration of 1.5% and a standardized OD_600_ of 0.2. Flow cytometry was performed using the BD LSRFortessa Cell Analyzer (BD Sciences, USA) equipped with an HTS autosampler (BD Sciences, USA) measuring sfGFP signals with a 488 nm laser and 530/30 nm detector and mKate with a 561 nm laser and 610/20 nm detector. Voltages for detecting forward scatter, side scatter, sfGFP and mKate were adjusted to 420, 270, 460 and 530, respectively. To reduce background debris, thresholds for forward and side scatter were set to 5,000, while 20,000 events were recorded.

### Induction assays

Overnight culture grown in absence of inducer was used to inoculated to media with various concentrations of cym and van in 96-well plates. The normalized steady-state fluorescence at late growth phase (Fss) averaged oved a time window of six to twelve hours was used as response variable of induction curves. In R (v4.3.1), Hill coefficient (*n*), activation coefficient (*K*) and max steady-state fluorescence output (β) was estimated by fitting the Hill function (1) using non-linear least-square regression with the “nls” function from the stats base R package. For parameter *C*, representing basal fluorescence output at 0 inducer concentration, empirical value was used.

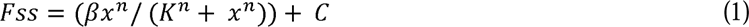

Where x is cym (mM) or van (mM) inducer concentration.

### Statistical analysis

All statistical analysis was done in R. Toggle switch performance metrics and growth metrics were unit-scaled prior to downstream analysis. Hierarchal clustering was done using “hclust” function (distance = “Euclidean”, method = “complete”) from base stats (v.3.6.2) package. Principal Component Analysis and Procrustes Superimposition analysis was done using the Vegan (v.2.6-4) package with functions “rda” and “protest” respectively. The *M*^2^ statistic from Procrustes Superimposition analysis (scale = TRUE, symmetric = TRUE) was tested for significance by a permutation approach (n = Inf, maximum number of iterations). Briefly, observations in one matrix are randomly reordered while maintaining the covariance structure within the matrix and a test statistic is calculated and recorded enough times to obtain a sizeable null distribution. A *P-*value for each statistic is then calculated, representing the probability of obtaining a statistic with a value equal to or more extreme of the experimental value.

## REFERENCES

1. Church, G. M., Elowitz, M. B., Smolke, C. D., Voigt, C. A. & Weiss, R. Realizing the potential of synthetic biology. Nat Rev Mol Cell Biol 15, 289–294 (2014).

2. Yan, X., Liu, X., Zhao, C. & Chen, G.-Q. Applications of synthetic biology in medical and pharmaceutical fields. Signal Transduct Target Ther 8, 199 (2023).

3. Symons, J. et al. Engineering biology and climate change mitigation: Policy considerations. Nat Commun 15, 2669 (2024).

4. Rodrigues, R. C., Pereira, H. S., Senra, R. L., Ribon, A. de O. B. & Mendes, T. A. de O. Understanding the emerging potential of synthetic biology for food science: Achievements, applications and safety considerations. Food Chemistry Advances 3, 100476 (2023).

5. Müller, K. M. & Arndt, K. M. Standardization in Synthetic Biology. in Synthetic Gene Networks: Methods and Protocols (eds. Weber, W. & Fussenegger, M.) 23–43 (Humana Press, Totowa, NJ, 2012). doi:10.1007/978-1-61779-412-4_2.

6. de Lorenzo, V. Evolutionary tinkering vs. rational engineering in the times of synthetic biology. Life Sciences, Society and Policy 14, 18 (2018).

7. Naseri, G. & Koffas, M. A. G. Application of combinatorial optimization strategies in synthetic biology. Nat Commun 11, 2446 (2020).

8. Mehrotra, R., Renganaath, K., Kanodia, H., Loake, G. J. & Mehrotra, S. Towards combinatorial transcriptional engineering. Biotechnology Advances 35, 390–405 (2017).

9. Catanach, T. A., McCardell, R., Baetica, A.-A. & Murray, R. M. Context Dependence of Biological Circuits. http://biorxiv.org/lookup/doi/10.1101/360040 (2018) doi:10.1101/360040.

10. Cardinale, S. & Arkin, A. P. Contextualizing context for synthetic biology--identifying causes of failure of synthetic biological systems. Biotechnol J 7, 856–866 (2012).

11. Adams, B. L. The Next Generation of Synthetic Biology Chassis: Moving Synthetic Biology from the Laboratory to the Field. ACS Synth. Biol. 5, 1328–1330 (2016).

12. Lee, H.-M. et al. Construction of a tunable promoter library to optimize gene expression in Methylomonas sp. DH-1, a methanotroph, and its application to cadaverine production. Biotechnology for Biofuels 14, 228 (2021).

13. Liebal, U. W. et al. Insight to Gene Expression From Promoter Libraries With the Machine Learning Workflow Exp2Ipynb. Front. Bioinform. 1, (2021).

14. Yuan, J. & Ching, C. B. Combinatorial Assembly of Large Biochemical Pathways into Yeast Chromosomes for Improved Production of Value-added Compounds. ACS Synth. Biol. 4, 23–31 (2015).

15. Ajikumar, P. K. et al. Isoprenoid Pathway Optimization for Taxol Precursor Overproduction in Escherichia coli. Science 330, 70–74 (2010).

16. Xu, P. et al. Modular optimization of multi-gene pathways for fatty acids production in E. coli. Nat Commun 4, 1409 (2013).

17. Duan, Y. et al. Deciphering the Rules of Ribosome Binding Site Differentiation in Context Dependence. ACS Synth. Biol. 11, 2726–2740 (2022).

18. Jeschek, M., Gerngross, D. & Panke, S. Rationally reduced libraries for combinatorial pathway optimization minimizing experimental effort. Nat Commun 7, 11163 (2016).

19. Smanski, M. J. et al. Functional optimization of gene clusters by combinatorial design and assembly. Nat Biotechnol 32, 1241–1249 (2014).

20. Zhang, S., Zhao, X., Tao, Y. & Lou, C. A novel approach for metabolic pathway optimization: Oligo-linker mediated assembly (OLMA) method. Journal of Biological Engineering 9, 23 (2015).

21. Feng, J. et al. Construction of cell factory through combinatorial metabolic engineering for efficient production of itaconic acid. Microbial Cell Factories 21, 275 (2022).

22. Wang, R. et al. Design and Characterization of Biosensors for the Screening of Modular Assembled Naringenin Biosynthetic Library in Saccharomyces cerevisiae. ACS Synth. Biol. 8, 2121–2130 (2019).

23. Juárez, J. F., Lecube-Azpeitia, B., Brown, S. L., Johnston, C. D. & Church, G. M. Biosensor libraries harness large classes of binding domains for construction of allosteric transcriptional regulators. Nat Commun 9, 3101 (2018).

24. Oesterle, S., Gerngross, D., Schmitt, S., Roberts, T. M. & Panke, S. Efficient engineering of chromosomal ribosome binding site libraries in mismatch repair proficient Escherichia coli. Sci Rep 7, 12327 (2017).

25. Zelcbuch, L. et al. Spanning high-dimensional expression space using ribosome-binding site combinatorics. Nucleic Acids Research 41, e98 (2013).

26. Ding, N. et al. Programmable cross-ribosome-binding sites to fine-tune the dynamic range of transcription factor-based biosensor. Nucleic Acids Res 48, 10602–10613 (2020).

27. Omotajo, D., Tate, T., Cho, H. & Choudhary, M. Distribution and diversity of ribosome binding sites in prokaryotic genomes. BMC Genomics 16, 604 (2015).

28. Failmezger, J., Ludwig, J., Nieß, A. & Siemann-Herzberg, M. Quantifying ribosome dynamics in Escherichia coli using fluorescence. FEMS Microbiology Letters 364, fnx055 (2017).

29. Salis, H. M., Mirsky, E. A. & Voigt, C. A. Automated Design of Synthetic Ribosome Binding Sites to Precisely Control Protein Expression. Nat Biotechnol 27, 946–950 (2009).

30. Roots, C. T., Lukasiewicz, A. & Barrick, J. E. OSTIR: open source translation initiation rate prediction. J Open Source Softw 6, 3362 (2021).

31. Calero, P. & Nikel, P. I. Chasing bacterial *chassis* for metabolic engineering: a perspective review from classical to non-traditional microorganisms. Microb. Biotechnol. 12, 98–124 (2019).

32. Smanski, M. J. et al. Synthetic biology to access and expand nature’s chemical diversity. Nat Rev Microbiol 14, 135–149 (2016).

33. Brooks, S. M. & Alper, H. S. Applications, challenges, and needs for employing synthetic biology beyond the lab. Nat Commun 12, 1390 (2021).

34. Khan, N., Yeung, E., Farris, Y., Fansler, S. J. & Bernstein, H. C. A broad-host-range event detector: expanding and quantifying performance between Escherichia coli and Pseudomonas species. Synthetic Biology 5, ysaa002 (2020).

35. Chan, D. T. C., Baldwin, G. S. & Bernstein, H. C. Revealing the Host-Dependent Nature of an Engineered Genetic Inverter in Concordance with Physiology. BioDesign Research 5, 0016 (2023).

36. Qian, Y., Huang, H.-H., Jiménez, J. I. & Del Vecchio, D. Resource Competition Shapes the Response of Genetic Circuits. ACS Synth Biol 6, 1263–1272 (2017).

37. Hartline, C. J. & Zhang, F. The Growth Dependent Design Constraints of Transcription-Factor-Based Metabolite Biosensors. ACS Synth. Biol. (2022) doi:10.1021/acssynbio.2c00143.

38. Müller, I. E. et al. Gene networks that compensate for crosstalk with crosstalk. Nat Commun 10, 4028 (2019).

39. Tas, H., Grozinger, L., Stoof, R., de Lorenzo, V. & Goñi-Moreno, Á. Contextual dependencies expand the re-usability of genetic inverters. Nat Commun 12, 355 (2021).

40. Ke, J. et al. Development of platforms for functional characterization and production of phenazines using a multi-chassis approach via CRAGE. Metabolic Engineering 69, 188–197 (2022).

41. Kim, J. et al. Properties of alternative microbial hosts used in synthetic biology: towards the design of a modular chassis. Essays Biochem 60, 303–313 (2016).

42. Kittleson, J. T., Wu, G. C. & Anderson, J. C. Successes and failures in modular genetic engineering. Current Opinion in Chemical Biology 16, 329–336 (2012).

43. Nikolados, E.-M., Weiße, A. Y., Ceroni, F. & Oyarzún, D. A. Growth Defects and Loss-of-Function in Synthetic Gene Circuits. ACS Synth. Biol. 8, 1231–1240 (2019).

44. Storch, M., Haines, M. C. & Baldwin, G. S. DNA-BOT: A low-cost, automated DNA assembly platform for synthetic biology. bioRxiv 832139 (2019) doi:10.1101/832139.

45. Storch, M. et al. BASIC: A New Biopart Assembly Standard for Idempotent Cloning Provides Accurate, Single-Tier DNA Assembly for Synthetic Biology. ACS Synth Biol 4, 781–787 (2015).

46. Haines, M. C. et al. basicsynbio and the BASIC SEVA collection: software and vectors for an established DNA assembly method. Synthetic Biology 7, ysac023 (2022).

47. Sridhar, S., Ajo-Franklin, C. M. & Masiello, C. A. A Framework for the Systematic Selection of Biosensor Chassis for Environmental Synthetic Biology. ACS Synth. Biol. 11, 2909–2916 (2022).

48. Guan, Y. et al. Mitigating Host Burden of Genetic Circuits by Engineering Autonegatively Regulated Parts and Improving Functional Prediction. ACS Synth. Biol. 11, 2361–2371 (2022).

49. Di Blasi, R. et al. Resource-aware construct design in mammalian cells. Nat Commun 14, 3576 (2023).

50. Cook, T. B. et al. Genetic tools for reliable gene expression and recombineering in Pseudomonas putida. J Ind Microbiol Biotechnol 45, 517–527 (2018).

51. Nikel, P. I. & de Lorenzo, V. Robustness of *Pseudomonas putida* KT2440 as a host for ethanol biosynthesis. New Biotechnology 31, 562–571 (2014).

52. Kaczmarczyk, A., Vorholt, J. A. & Francez-Charlot, A. Cumate-Inducible Gene Expression System for Sphingomonads and Other Alphaproteobacteria. Appl Environ Microbiol 79, 6795–6802 (2013).

53. Seo, S.-O. & Schmidt-Dannert, C. Development of a synthetic cumate-inducible gene expression system for Bacillus. Appl Microbiol Biotechnol 103, 303–313 (2019).

54. Kaczmarczyk, A., Vorholt, J. A. & Francez-Charlot, A. Synthetic vanillate-regulated promoter for graded gene expression in Sphingomonas. Sci Rep 4, 6453 (2014).

55. Thanbichler, M., Iniesta, A. A. & Shapiro, L. A comprehensive set of plasmids for vanillate- and xylose-inducible gene expression in Caulobacter crescentus. Nucleic Acids Res 35, e137 (2007).

56. Bienick, M. S. et al. The Interrelationship between Promoter Strength, Gene Expression, and Growth Rate. PLoS One 9, e109105 (2014).

57. Pougach, K. et al. Duplication of a promiscuous transcription factor drives the emergence of a new regulatory network. Nat Commun 5, 4868 (2014).

58. Chan, D. T. C. & Bernstein, H. C. Pangenomic Landscapes Shape Performances of a Synthetic Genetic Circuit Across Stutzerimonas Species. 2024.02.15.580380 Preprint at 10.1101/2024.02.15.580380 (2024).

59. Klumpp, S., Zhang, Z. & Hwa, T. Growth-rate dependent global effects on gene expression in bacteria. Cell 139, 1366 (2009).

60. Carbonell, P., Radivojevic, T. & García Martín, H. Opportunities at the Intersection of Synthetic Biology, Machine Learning, and Automation. ACS Synth. Biol. 8, 1474–1477 (2019).

61. Yáñez Feliú, G., et al. Flapjack: Data Management and Analysis for Genetic Circuit Characterization. ACS Synth. Biol. 10, 183–191 (2021).

62. Boada, Y. et al. Characterization of Gene Circuit Parts Based on Multiobjective Optimization by Using Standard Calibrated Measurements. ChemBioChem 20, 2653–2665 (2019).

63. Stone, A., Youssef, A., Rijal, S., Zhang, R. & Tian, X.-J. Context-dependent redesign of robust synthetic gene circuits. Trends in Biotechnology (2024) doi:10.1016/j.tibtech.2024.01.003.

64. LeBlanc, N. & Charles, T. C. Bacterial genome reductions: Tools, applications, and challenges. Front. Genome Ed. 4, (2022).

65. Darlington, A. P. S., Kim, J., Jiménez, J. I. & Bates, D. G. Dynamic allocation of orthogonal ribosomes facilitates uncoupling of co-expressed genes. Nature Communications 9, 695 (2018).

66. Costello, A. & Badran, A. H. Synthetic Biological Circuits Within an Orthogonal Central Dogma. Trends Biotechnol 39, 59–71 (2021).

67. Meyer, A. J., Ellefson, J. W. & Ellington, A. D. Directed Evolution of a Panel of Orthogonal T7 RNA Polymerase Variants for in Vivo or in Vitro Synthetic Circuitry. ACS Synth. Biol. 4, 1070–1076 (2015).

68. Ostrov, N. et al. Synthetic genomes with altered genetic codes. Current Opinion in Systems Biology 24, 32–40 (2020).

69. Martínez-García, E. & de Lorenzo, V. The quest for the minimal bacterial genome. Current Opinion in Biotechnology 42, 216–224 (2016).

70. Hutchison, C. A. et al. Design and synthesis of a minimal bacterial genome. Science 351, aad6253 (2016).

71. Sung, B. H., Choe, D., Kim, S. C. & Cho, B.-K. Construction of a minimal genome as a chassis for synthetic biology. Essays in Biochemistry 60, 337–346 (2016).

72. Kwon, K. K., Lee, J., Kim, H., Lee, D.-H. & Lee, S.-G. Advancing high-throughput screening systems for synthetic biology and biofoundry. Current Opinion in Systems Biology 37, 100487 (2024).

73. Ikeda, H., Shin-ya, K. & Omura, S. Genome mining of the Streptomyces avermitilis genome and development of genome-minimized hosts for heterologous expression of biosynthetic gene clusters. Journal of Industrial Microbiology and Biotechnology 41, 233–250 (2014).

74. Morimoto, T. et al. Enhanced Recombinant Protein Productivity by Genome Reduction in Bacillus subtilis. DNA Res 15, 73–81 (2008).

75. Wynands, B. et al. Streamlined Pseudomonas taiwanensis VLB120 Chassis Strains with Improved Bioprocess Features. ACS Synth. Biol. 8, 2036–2050 (2019).

76. Wang, G. et al. CRAGE enables rapid activation of biosynthetic gene clusters in undomesticated bacteria. Nat Microbiol 4, 2498–2510 (2019).

77. Bertile, F., Matallana-Surget, S., Tholey, A., Cristobal, S. & Armengaud, J. Diversifying the concept of model organisms in the age of -omics. Commun Biol 6, 1–4 (2023).

78. Duffy, M. A. et al. Model Systems in Ecology, Evolution, and Behavior: A Call for Diversity in Our Model Systems and Discipline. The American Naturalist 198, 53–68 (2021).

79. Fatma, Z., Schultz, J. C. & Zhao, H. Recent advances in domesticating non-model microorganisms. Biotechnology Progress 36, e3008 (2020).

80. Liu, H. & Deutschbauer, A. M. Rapidly moving new bacteria to model-organism status. Current Opinion in Biotechnology 51, 116–122 (2018).

81. Sambrook, J. & Russell, D. W. The inoue method for preparation and transformation of competent e. Coli: ‘ultra-competent’ cells. CSH Protoc 2006, pdb.prot3944 (2006).

82. Shine, J. & Dalgarno, L. The 3’-terminal sequence of Escherichia coli 16S ribosomal RNA: complementarity to nonsense triplets and ribosome binding sites. Proc. Natl. Acad. Sci. U.S.A. 71, 1342–1346 (1974).

83. Lorenz, R. et al. ViennaRNA Package 2.0. Algorithms for Molecular Biology 6, 26 (2011).

84. Zwietering, M. H., Jongenburger, I., Rombouts, F. M. & van ‘t Riet, K. Modeling of the Bacterial Growth Curve. Appl Environ Microbiol 56, 1875–1881 (1990).

