## Supplementary Figures, Tables and Other for "Fine Tuning Genetic Circuits via Host Context and RBS Modulation"

**Supplementary Table S1 – S2**

**Supplementary Material S1**

Dennis Tin Chat Chan^1^, Lena Winter^1^, Johan Bjerg^1^, Stina Krsmanovic^1^, Geoff S. Baldwin^3,4^, Hans C. Bernstein^1,2^*

^1^Faculty of Biosciences, Fisheries and Economics, UiT - The Arctic University of Norway, 9019, Tromsø, Norway

^2^The Arctic Centre for Sustainable Energy, UiT - The Arctic University of Norway, 9019, Tromsø, Norway

^3^Department of Life Sciences, Imperial College London, South Kensington, London SW7 2AZ, UK

^4^Imperial College Centre for Synthetic Biology, Imperial College London, South Kensington, London SW7 2AZ, UK

Keywords:

Synthetic Biology, Toggle Switch, Chassis-Effect, Context Dependence, Biodesign, RBS, Genetic Circuit, Modularity, Stutzerimonas, Non-model, Host Organism, Host-Circuit Interaction.

### Supplementary Figures

#### Supplementary Figure S1

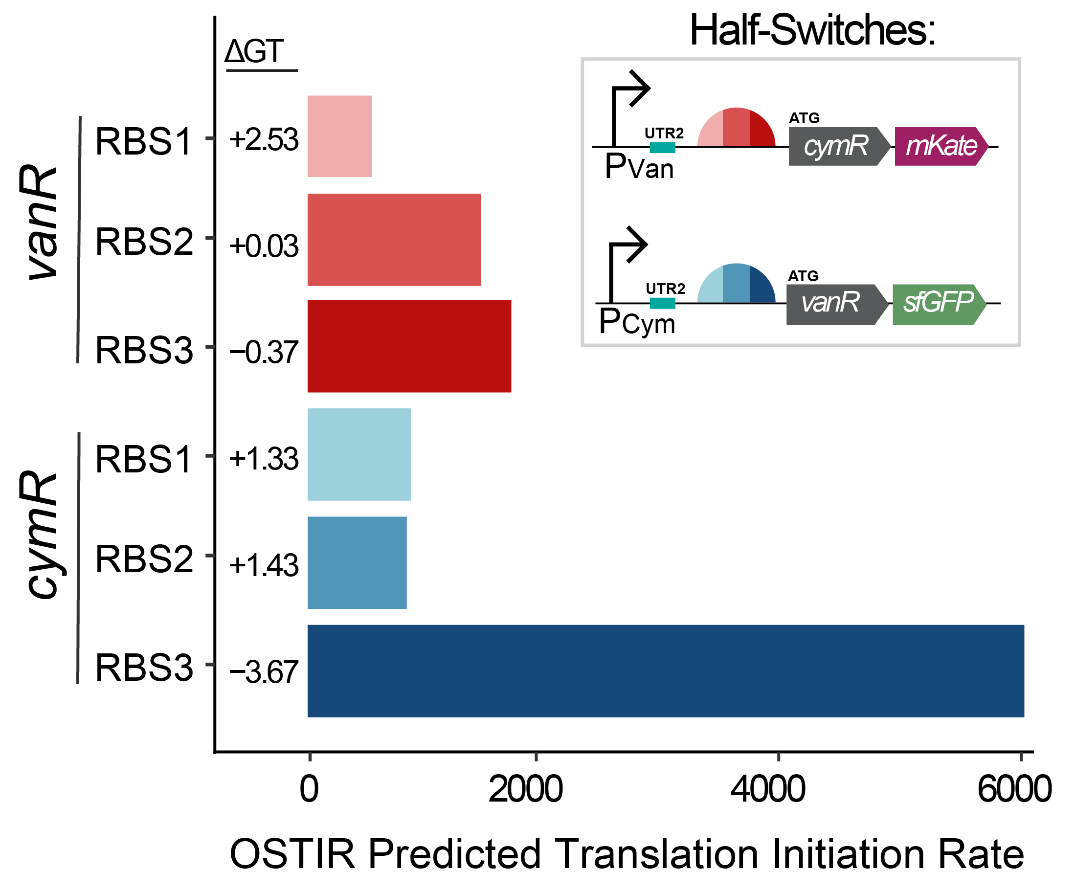

**Supplementary Figure S1. Estimated translation initiation rate is context dependent.** Inferred translation initiation rate of each RBS context regulating for *cymR* and *vanR* genes within the toggle switches by OSTIR. ΔGT: change in Gibb’s free energy associated with ribosome binding to the RBS part.

#### Supplementary Figure S2

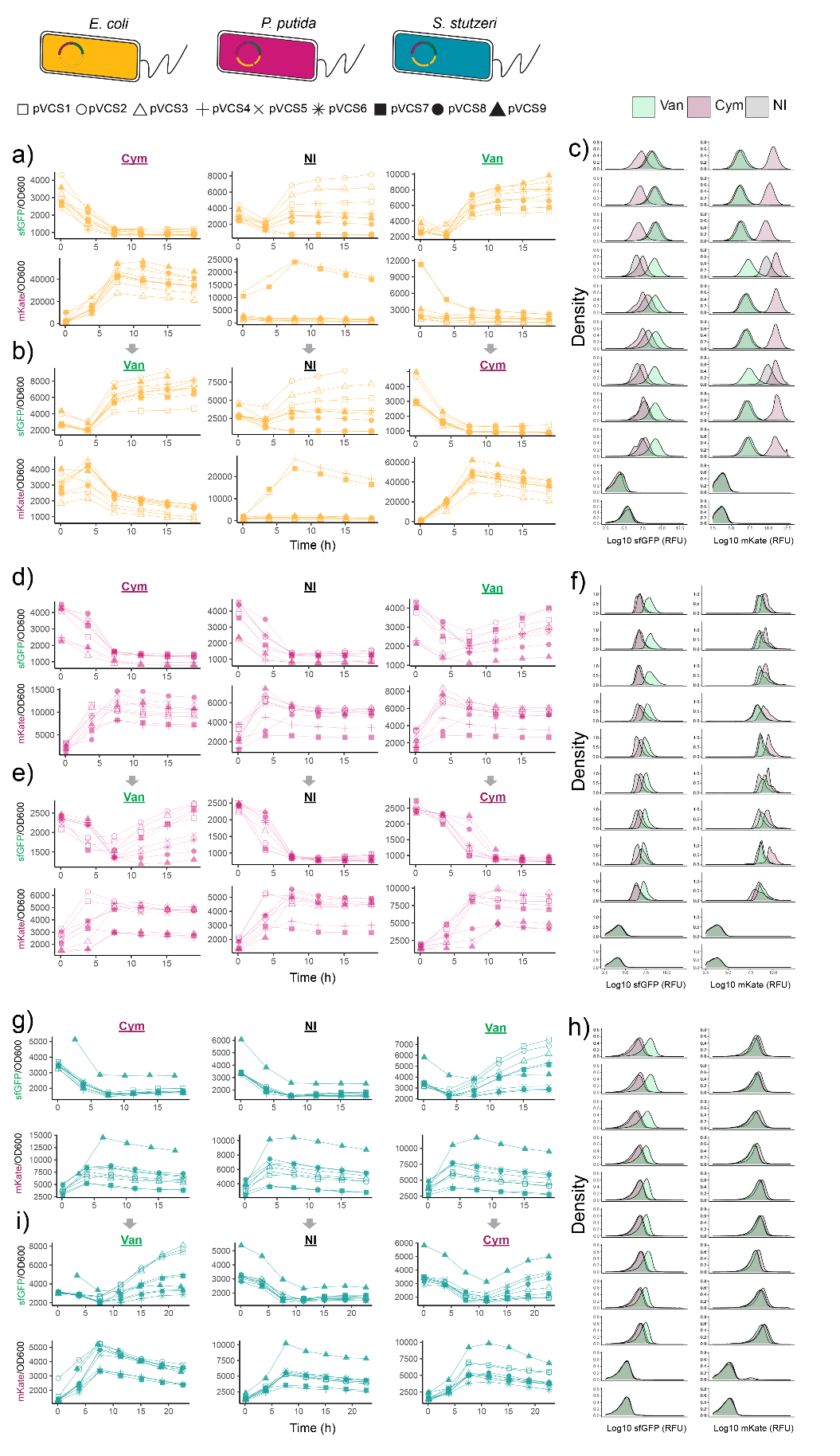

**Supplementary Figure S2. Chassis-effect revealed through toggle switch fluorescence dynamics.** Normalized fluorescence dynamics of toggled cells and fluorescence intensity distribution of late-phase cells for a-c) *Escherichia coli* DH5α, d-e) *Pseudomonas putida* KT2440 and g-i) *Stutzerimonas stutzeri* CCUG 11256*.* Cym: cumate; van: vanillate; NI: no inducer.

#### Supplementary Figure S3

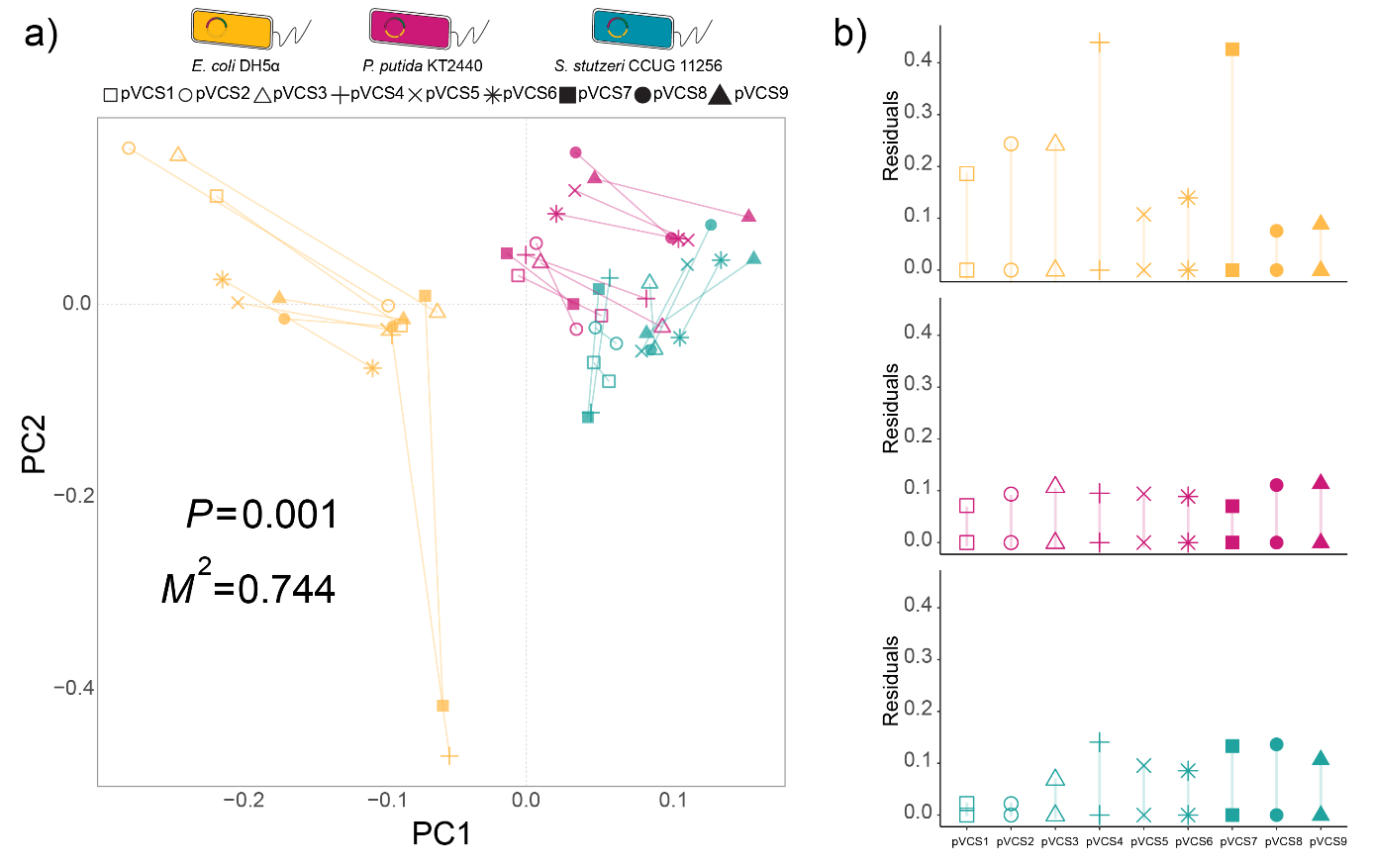

**Supplementary Figure S3. Differences in growth between toggle switch variants significantly correlated with differences in toggle switch performance.** a) PCA-based Procrustes Superimposition analysis of differential growth dynamics and differential toggle switch performance (toggle assay and induction metrics). Connecting lines indicate vector residuals. *P*: p-value. *M*^2^: Gower statistic, the sum of squared vector residuals. b) Bar plot of residual vector between points.

### Supplementary Table

#### Supplementary Table S1

**Supplementary Table 1.** Species used in this study.

| **Name** | **Culture Collection Acc.** | **NCBI Assembly Acc.** | **Genotype** | **Reference** |
| --- | --- | --- | --- | --- |
| *Escherichia coli* DH5α | DSM 6897 | GCF_002899475.1 | WT | ^1^ |
| *Pseudomonas putida* KT2440 | DSM 6125 | GCF_000007565.2 | WT | ^2^ |
| *Stutzerimonas stutzeri* CCUG11256 | CCUG11256 | GCF_000219605.1 | WT | ^3^ |
| *Escherichia coli* DH5α | DSM 6897 | NA | pVCS1 | This Study |
| *Escherichia coli* DH5α | DSM 6897 | NA | pVCS2 | This Study |
| *Escherichia coli* DH5α | DSM 6897 | NA | pVCS3 | This Study |
| *Escherichia coli* DH5α | DSM 6897 | NA | pVCS4 | This Study |
| *Escherichia coli* DH5α | DSM 6897 | NA | pVCS5 | This Study |
| *Escherichia coli* DH5α | DSM 6897 | NA | pVCS6 | This Study |
| *Escherichia coli* DH5α | DSM 6897 | NA | pVCS7 | This Study |
| *Escherichia coli* DH5α | DSM 6897 | NA | pVCS8 | This Study |
| *Escherichia coli* DH5α | DSM 6897 | NA | pVCS9 | This Study |
| *Pseudomonas putida* KT2440 | DSM 6125 | NA | pVCS1 | This Study |
| *Pseudomonas putida* KT2440 | DSM 6125 | NA | pVCS2 | This Study |
| *Pseudomonas putida* KT2440 | DSM 6125 | NA | pVCS3 | This Study |
| *Pseudomonas putida* KT2440 | DSM 6125 | NA | pVCS4 | This Study |
| *Pseudomonas putida* KT2440 | DSM 6125 | NA | pVCS5 | This Study |
| *Pseudomonas putida* KT2440 | DSM 6125 | NA | pVCS6 | This Study |
| *Pseudomonas putida* KT2440 | DSM 6125 | NA | pVCS7 | This Study |
| *Pseudomonas putida* KT2440 | DSM 6125 | NA | pVCS8 | This Study |
| *Pseudomonas putida* KT2440 | DSM 6125 | NA | pVCS9 | This Study |
| *Stutzerimonas stutzeri* CCUG11256 | CCUG11256 | NA | pVCS1 | This Study |
| *Stutzerimonas stutzeri* CCUG11256 | CCUG11256 | NA | pVCS2 | This Study |
| *Stutzerimonas stutzeri* CCUG11256 | CCUG11256 | NA | pVCS3 | This Study |
| *Stutzerimonas stutzeri* CCUG11256 | CCUG11256 | NA | pVCS4 | This Study |
| *Stutzerimonas stutzeri* CCUG11256 | CCUG11256 | NA | pVCS5 | This Study |
| *Stutzerimonas stutzeri* CCUG11256 | CCUG11256 | NA | pVCS6 | This Study |
| *Stutzerimonas stutzeri* CCUG11256 | CCUG11256 | NA | pVCS7 | This Study |
| *Stutzerimonas stutzeri* CCUG11256 | CCUG11256 | NA | pVCS8 | This Study |
| *Stutzerimonas stutzeri* CCUG11256 | CCUG11256 | NA | pVCS9 | This Study |

#### Supplementary Table S2

**Supplementary Table 2.** DNA sequences of parts used in this study.

| **Part** | **Sequence** |
| --- | --- |
| PCym | CTCGGTACCAAATTCCAGAAAAGAGACGCTTTCGAGCGTCTTTTTTCGTTTTGGTCCGTGCCTACTCTGGAAAATCTAACAAACAGACAATCTGGTCTGTTTGTATTATGGAAAATTTTTCTGTATAATAGATTCAACAAACAGACAATCTGGTCTGTTTGTATTATAGCGCTCAACGGGTGTGCTTCCCGTTCTGATGAGTCCGTGAGGACGAAAGCGCCTCTACAAATAATTTTGTTTAA |
| PVan | GACGAACAATAAGGCCTCCCTAACGGGGGGCCTTTTTTATTGATAACAAAAGTGCCTACTCTGGAAAATCTATTGGATCCAATTGACAGCTAGCTCAGTCCTAGGTACCATTGGATCCAATAGTAGTCACCGGCTGTGCTTGCCGGTCTGATGAGCCTGTGAAGGCGAAACTACCTCTACAAATAATTTTGTTTAA |
| *cymR* | ATGAGCCCGAAACGTCGTACCCAGGCAGAACGTGCAATGGAAACCCAGGGTAAACTGATTGCAGCAGCACTGGGTGTTCTGCGTGAAAAAGGTTATGCAGGTTTTCGTATTGCAGATGTTCCGGGTGCAGCCGGTGTTAGCCGTGGTGCACAGAGCCATCATTTTCCGACCAAACTGGAACTGCTGCTGGCAACCTTTGAATGGCTGTATGAGCAGATTACCGAACGTAGCCGTGCACGTCTGGCAAAACTGAAACCGGAAGATGATGTTATTCAGCAGATGCTGGATGATGCAGCAGAATTTTTTCTGGATGATGATTTTAGCATCGGCCTGGATCTGATTGTTGCAGCAGATCGTGATCCGGCACTGCGTGAAGGTATTCAGCGTACCGTTGAACGTAATCGTTTTGTTGTTGAAGATATGTGGCTGGGTGTGCTGGTGAGCCGTGGTCTGAGCCGTGATGATGCCGAAGATATTCTGTGGCTGATTTTTAACAGCGTTCGTGGTCTGGTAGTTCGTAGCCTGTGGCAGAAAGATAAAGAACGTTTTGAACGTGTGCGTAATAGCACCCTGGAAATTGCACGTGAACGTTATGCAAAATTCAAACGTTGA |
| *vanR* | ATGGACATGCCTCGTATTAAACCGGGTCAGCGTGTTATGATGGCACTGCGTAAAATGATTGCAAGCGGTGAAATCAAAAGTGGTGAACGTATTGCAGAAATTCCGACCGCAGCAGCACTGGGTGTTAGCCGTATGCCGGTTCGTATCGCACTGCGTTCACTGGAACAAGAAGGTCTGGTTGTTCGTCTGGGTGCACGTGGTTATGCAGCCCGTGGTGTTAGCAGCGATCAGATTCGTGATGCAATTGAAGTTCGTGGTGTTCTGGAAGGTTTTGCAGCACGTCGTCTGGCAGAACGTGGTATGACCGCAGAAACCCATGCACGTTTTGTTGTACTGATTGCAGAAGGTGAAGCACTGTTTGCAGCCGGTCGCCTGAATGGTGAAGATCTGGATCGTTATGCCGCATATAATCAGGCATTTCATGATACCCTGGTTAGCGCAGCAGGTAATGGTGCAGTTGAAAGCGCACTGGCACGTAATGGTTTTGAACCGTTTGCAGCAGCCGGTGCACTGGCCCTGGATCTGATGGACCTGTCTGCCGAATATGAACATCTGCTGGCAGCACATCGTCAGCATCAGGCAGTTCTGGATGCAGTTAGCTGTGGTGATGCCGAAGGTGCAGAACGTATTATGCGTGATCATGCACTGGCAGCAATTCGTAATGCAAAAGTTTTTGAAGCAGCAGCAAGCGCAGGCGCACCGCTGGGTGCAGCATGGTCAATTCGTGCAGATTGA |
| *sfGFP* | ATGCGTAAAGGCGAAGAGCTGTTCACTGGTGTCGTCCCTATTCTGGTGGAACTGGATGGTGATGTCAACGGTCATAAGTTTTCCGTGCGTGGCGAGGGTGAAGGTGACGCAACTAATGGTAAACTGACGCTGAAGTTCATCTGTACTACTGGTAAACTGCCGGTACCTTGGCCGACTCTGGTAACGACGCTGACTTATGGTGTTCAGTGCTTTGCTCGTTATCCGGACCATATGAAGCAGCATGACTTCTTCAAGTCCGCCATGCCGGAAGGCTATGTGCAGGAACGCACGATTTCCTTTAAGGATGACGGCACGTACAAAACGCGTGCGGAAGTGAAATTTGAAGGCGATACTCTGGTAAACCGCATTGAGCTGAAAGGCATTGACTTTAAAGAAGACGGCAATATCCTGGGCCATAAGCTGGAATACAATTTTAACAGCCACAATGTTTACATCACCGCCGATAAACAAAAAAATGGCACTAAAGCGAATTTTAAAATTCGCCACAACGTGGAGGATGGCAGCGTGCAGCTGGCTGATCACTACCAGCAAAACACTCCAATCGGTGATGGTCCTGTTCTGCTGCCAGACAATCACTATCTGAGCACGCAAAGCGTTCTGTCTAAAGATCCGAACGAGAAACGCGATCATATGGTTCTGCTGGAGTTCGTAACCGCAGCGGGCATCACGCATGGTATGGATGAACTGTAC |
| *mKate* | ATGTCAGAATTAATTAAAGAAAATATGCACATGAAATTATATATGGAAGGTACTGTCAACAATCATCATTTCAAATGCACATCCGAAGGTGAAGGTAAACCATATGAAGGCACACAAACAATGCGCATCAAAGCAGTTGAAGGTGGACCCCTGCCCTTTGCGTTTGACATTCTCGCAACGAGCTTTATGTACGGGTCTAAAACTTTTATCAATCACACCCAAGGCATTCCTGACTTTTTTAAACAGTCCTTTCCTGAAGGCTTTACCTGGGAACGTGTAACAACTTATGAAGATGGCGGTGTACTTACAGCAACTCAAGATACGAGTTTACAAGATGGCTGTCTGATTTACAATGTTAAAATCCGTGGCGTAAATTTCCCGAGTAACGGACCCGTAATGCAAAAAAAAACTCTTGGTTGGGAAGCATCAACAGAAACCTTATATCCTGCGGACGGTGGCTTAGAAGGACGCGCAGACATGGCACTGAAATTAGTTGGAGGCGGTCATTTAATCTGCAACCTGAAAACAACCTATCGTTCCAAAAAACCCGCTAAAAACCTTAAAATGCCTGGAGTATACTATGTTGATCGTCGCTTAGAACGTATTAAAGAAGCTGATAAAGAAACCTACGTTGAACAACATGAAGTAGCCGTAGCCCGTTATTGTGACCTTCCGTCGAAATTAGGACATCGTTGA |
| UTR1 | TTGAACACCGTCTCAGGTAAGTATCAGTTGTA |
| UTR2 | TGTTACTATTGGCTGAGATAAGGGTAGCAGAA |
| UTR3 | GTATCTCGTGGTCTGACGGTAAAATCTATTGT |
| RBS1 | ATCACACAGGACTA |
| RBS2 | AAAGAGGGGAAATA |
| RBS3 | AAAGAGGAGAAATA |

### Supplementary Material

#### Supplementary Material S1

**DNA-BOT Application Protocol Description**

DNA-BOT was used to generate the four scripts to program the OpenTrons2 pipetting robot to assemble our toggle switches via BASIC DNA assembly. DNA-BOT application was run with default settings and the protocol was followed as described by Storch et al. (2020)^4^ with the following differences:

In “2_purification_ot2_APIv2.8.py” script, the following lines were replaced with:

| **Line** | **Replaced syntax** |
| --- | --- |
| 125 | pipette.flow_rate.aspirate=PIPETTE_ASPIRATE_RATE |
| 126 | pipette.flow_rate.dispense=PIPETTE_DISPENSE_RATE |

The following settings for labware ID and parameters were changed to the indicated value:

| **Step 2 – Labware IDs** | **Value** |
| --- | --- |
| Opentrons magnetic module gen2 | magnetic_module_gen2 |
| Opentrons 4-in-1 tubes rack | opentrons_24_tuberack_eppendorf_1.5ml_safelock_snapcap |
| 96 well rigid PCR plate (clip and transformation steps) | armadillo_96_wellplate_200ul_pcr_full_skirt |
| 96 well rigid PCR plate (purification and assembly steps) | armadillo_96_wellplate_200ul_pcr_full_skirt |
| Agar plate (transformation step) | armadillo_96_wellplate_200ul_pcr_full_skirt |
| Reservoir plate 21 mL 12 channels | nest_12_reservoir_15ml |
| 96 deep well plate 2 mL wells | thermoscientificnunc_96_wellplate_2000ul |

| **Step 3 - Parameters** | **Value** |
| --- | --- |
| Magnetic module height (mm) | 11 |
| Settling time (min) | 5 |

### References

1.	Chen, J., Li, Y., Zhang, K. & Wang, H. Whole-Genome Sequence of Phage-Resistant Strain Escherichia coli DH5α. *Genome Announcements* **6**, 10.1128/genomea.00097-18 (2018).

2.	Martin-Pascual, M. *et al.* A navigation guide of synthetic biology tools for Pseudomonas putida. *Biotechnology Advances* **49**, 107732 (2021).

3.	Gomila, M., Mulet, M., García-Valdés, E. & Lalucat, J. Genome-Based Taxonomy of the Genus Stutzerimonas and Proposal of S. frequens sp. nov. and S. degradans sp. nov. and Emended Descriptions of S. perfectomarina and S. chloritidismutans. *Microorganisms* **10**, 1363 (2022).

4.	Storch, M., Haines, M. C. & Baldwin, G. S. DNA-BOT: a low-cost, automated DNA assembly platform for synthetic biology. *Synthetic Biology* **5**, ysaa010 (2020).
